# The role of ADAR editing and nonsense-mediated decay in Parkinson’s Disease

**DOI:** 10.1101/2024.05.17.594716

**Authors:** Heather Milliken Mercer, Aiswarya Mukundan Nair, Ayesha Tariq, Helen Piontkivska

**Affiliations:** University of Mount Union; Kent State University

**Keywords:** Parkinson’s Disease, ADAR editing, RNA editing, nonsense-mediated decay

## Abstract

Parkinson’s Disease (PD) is a multifactorial disease with heterogenous phenotypes that vary across individuals, as well as by age and sex. Therefore, it is likely that multiple interacting factors, such as environmental influences and aging, as well as genetic factors, including dynamic RNA (ADAR, Adenosine Deaminases Acting on RNA) editing, may play a role in PD pathology. In this analysis of 317 transcriptomes of healthy controls, PD and prodromal patients aged 65 years or older, from Parkinson’s Project Markers Initiative dataset, we observe differences in ADAR expression, number of putative ADAR edits, editing index, and the number of high and moderate impact edits between control groups and diseased samples, particularly when ADAR editing is associated with nonsense-mediated decay (NMD). Likewise, differentially expressed genes between comparison groups were linked to NMD-related pathways. NMD is an important process in detecting deleterious nonsense sequences in mRNA transcripts and eliminating them from the cell. Thus, NMD regulation serves an important role in neurodevelopment, neural differentiation, and neural maturation. RNA misprocessing, which includes dysregulation of NMD, is known to play an important role in neurodegenerative diseases such as amyotrophic lateral sclerosis (ALS) and fronto-temporal dementia. Our results suggest that NMD may also be an important factor in PD physiology.

## 1 Introduction

The etiology of Parkinson’s Disease (PD) is multifactorial, including both genetic and environmental origins. Among the environmental factors, viral infections (influenza, Coxsackie, Japanese encephalitis virus {JEV}, and acquired immunodeficiency viruses) have been recognized as an independent risk for PD [1,2]. It has been further hypothesized that dysregulation of the innate and adaptive immune response is linked to the neuroinflammation present in PD progression as post-mortem, brain imaging, epidemiological, and animal studies demonstrating its involvement in neurodegeneration[3–9]. While multiple individual genes have been the focus of PD progression in recent studies[10–14], other genetic and environmental influences may also contribute to the physiological manifestation of PD. RNA editing by Adenosine Deaminases Acting on RNA (ADARs), which causes adenosine to inosine deamination, is the most common mechanism of post-transcriptional RNA editing in the nervous system and is integral to neurodevelopment and response to infection/inflammation[15–18]. PD is a complex disease with heterogenous phenotypes that vary across individuals, as well as by age and sex[19–21], therefore, it is likely that multiple interacting factors, such as the aging process[22,23] and hormones[24–26], in addition to genetic[11–14,27] and other confounding factors, including dynamic ADAR editing[14,28,29], may play a role in PD pathology.

## Background

While ∼15 million RNA editing sites have been observed in humans[30], only a portion of them are found in protein coding regions and lead to non-synonymous coding[28]. However, these substitutions are over-represented in transcripts known to function in neural tissues[31]. In a recent study of BA9 brain samples from a small cohort of PD patients, differential A-to-I editing was observed in 160 sites, with half of those edits occurring in genes encoding glutamate ionotropic receptors. Overall, Pozydyshev et al.[28] (2021) observed decreased editing in post-mortem PD brains as compared to controls. On the other hand, we recently [32] observed elevated levels of high and moderate impact protein-coding ADAR edits in skeletal muscle tissue in genes with known associations to PD in a small males-only cohort engaged in exercise. In a study examining post-mortem brain tissue for editing in microRNA (miRNA), Lu et al.[33], identified a number of miRNA editing sites that were edited at significantly different editing levels between PD and healthy controls (2022). Li et al. (2024) observed multiple ADAR editing sites that could be integral to PD physiology in a recent study of PD blood samples[34]. These findings suggest that changes in ADAR editing patterns may play an impactful role in the pathology of PD affecting various tissue types.

Nonsense-mediated decay (NMD) is an mRNA surveillance mechanism in eukaryotes, which largely serves to eliminate nonsense mutations[35,36] that arise from a variety of reasons, including ADAR editing, and function to regulate cellular processes such as stress responses[37]. In the nervous system, NMD moderates neurodevelopment, neurogenesis, neuronal differentiation[38,39], and synaptic plasticity[40,41]. NMD dysregulation has been implicated in intellectual disabilities[42], neurodevelopmental disorders[43,44], Amyotrophic Lateral Sclerosis (ALS), frontotemporal dementia, and PD[45,46]. A study of NMD events in the leukocytes of PD patients comparing samples taken before Deep Brain Stimulation (DBS) therapy and after DBS demonstrated that NMD events increased with disease progression, suggesting that NMD is an important regulator of the transcriptome, but also that brain changes transcend into peripheral systems[45]. Soreq et al. also noted that NMD and alternative splicing events, along with a reduction in tremors associated with PD, decreased for a time period following DBS[45]. NMD events are commonly associated with DNA repair genes and those that regulate alternative splicing. Collectively, Soreq et al.’s results suggest that NMD events may provide a target for potential PD treatment[45]. The targeting of NMD and NMD-related factors has also been suggested as a potential therapy for ALS and other neurodegenerative disorders[43].

Although PD pathology is most evident in the brain, crosstalk between the CNS and peripheral organ systems provides clues to PD pathophysiology including increasing evidence that circulating serum neurofilament light chains[47], small non-coding RNAs[48], and circulating cytokines[49] found in blood vary between PD and control samples. Additionally, altered cytosine-methylation, abnormal gene expression[50], and altered miRNAs[51] have been observed when comparing blood samples between PD patients and healthy individuals. In a recent study of RNA editing events in PD blood samples, the majority of edits occurred in protein-coding regions, non-exonic splicing regions, and Alu repeats, although a decreased overall editing frequency was observed when compared to controls[52]. The genes associated with the edits were enriched in neurodegeneration-related functions, with a subset of edits determined to change miRNA binding sites[52]. Additionally, a recent study of A to I editing in the transcriptomes of 380 PD patients and 178 Healthy Controls identified 17 ADAR edited sites that could contribute to increased genetic risk of PD through RNA editing[34]. While the 17 sites were associated with NCOR, KANSL, and BST genes in the Li et al.[34] study (2023), 57 additional sites were correlated with the progression of cognitive symptoms in PD cases[34].

To further confound the etiology of PD, phenotypes are not consistent between individual patients or within patients grouped by age or sex[53,54]. Historically, PD subtypes were characterized by one variable such as age or tremor[55], but have expanded to include a wider variety of symptoms, including sex[56]. Males are diagnosed with PD at a much higher rate than females[57,58], and females generally experience later onset yet faster disease progression than males[58,59]. It has been suggested that estrogen is neuroprotective and contributes to some of the discrepancies in onset and symptomology between the sexes[60]. Sex differences in PD progression have been well documented, including alterations in motor features by body region and postural scores[61], increased drooling[62,63], speech problems[62], more pronounced cognitive impairment, daytime sleepiness and swallowing in men[63,64], postural instability in women[65,66], and recently, Sandor et al. (2022) observed a significant association between sex and anxiety and depression[54].

The heterogeneity observed in PD cases has not been fully explained through genetic, age-related, or environmental associations[67], therefore we wanted to explore the associations between ADAR editing and the physiology of PD. Using blood-derived transcriptomes from 317 healthy controls, PD and prodromal patients aged 65 years or older, obtained from the Parkinson’s Progression Markers Initiative (PPMI)[68], we observed *that changes in ADAR editing accompany physiological changes in PD, resulting in dysfunctional/dysregulated proteins, and that these changes manifest differently across patients based on disease progression and sex, specifically in editing resulting in NMD and genes related to NMD physiology*.

## 2 Materials and Methods

In this analysis, the RNA-seq baseline data derived from blood samples of 179 PD, 64 healthy control, and 74 prodromal participants aged 65 years or older was obtained from the PPMI[68]. RNA-seq data used in the preparation of this analysis was obtained on October 31, 2021, from the PPMI database (www.ppmi-info.org/access-dataspecimens/download-data), RRID:SCR 006431, with an average of 114 M(illion) reads (mean reads = 116M paired reads/sample). The PD group consisted of 114 males and 65 females, the healthy control group consisted of 43 males and 21 females, and the prodromal group contained 25 females and 49 males, all 65 years of age or older. For all analyses, comparisons were made between healthy controls, PD, and prodromal samples in which both males and females are found, and between males and females within and between groups.

The Automated Isoform Diversity Detector (AIDD) pipeline[69] was used to infer ADAR editing events. Briefly, fastq files of individual patients were trimmed and aligned to the chosen human reference (GRCh37) using HISAT2[70]. Once aligned, the transcriptome was assembled using StringTie[71], to estimate levels of individual gene expression as Transcripts Per Kilobase Million (TPM). Putative ADAR editing events here are those A to G or U(T) to C (referred subsequently as T-to C) edits without Single Nucleotide Polymorphism Database annotation (dbSNP)[72] associations that have been assigned as “Pass” (filters) or have met “basic snp filter” criteria as designated by GATK variant calling pipeline[73] in AIDD[69]. SnpEff[74] was used to predicts the effects and functional impacts of predicted edited variants, as described below. Variants identified are putative ADAR editing events described as such based on known ADAR editing sites, but for simplicity will be further described throughout this study as simply “editing events” or “edits”.

A list of 801 unique gene IDs with suspected associations to PD were downloaded from NCBI Gene[75] (downloaded on November 4, 2023) and converted to Ensembl gene and transcript IDs, resulting in 901 gene and 7311 transcript IDs (S1 File) utilizing Ensembl Biomart[76]. The NCBI PD Gene List[75] (n=901) is one of the multiple GeneRIF (Gene Relevance into Function) lists, which are created and frequently updated to reflect current research through three primary methods: extraction from published literature by the National Library of Medicine staff, summary reports from HuGE Navigator[77], and user submissions from a Gene record[78]. The NCBI Gene List can be found in Supplementary File 1.

We used Reactome[79] to assess the pathways in which the NCBI PD genes were overrepresented utilizing two filters for stringency. The most stringent filter assessed only Reactome pathways with an FDR value < 0.05, while a less stringent filter included pathways with a p value < 0.05 with at least 50% of entities in the pathway.

SnpEff[74] annotations were collated by sample and patient group. Edited NCBI PD genes found within PD, healthy controls, and prodromal samples were compared between groups and analyzed for significant difference between groups (using Chi-square test). Only genes affected by high or moderate impact editing events occurring at a specific site in at least 25% of samples within a group were considered, similar to procedures followed by other studies[80–82] in order to reduce stochastic editing events. Within comparison groups, using the limit of genes edited in at least 25% of samples, resulted in edits by coordinate occurring in at least 6 healthy control females, 11 healthy control males, 17 PD females, 29 PD males, 7 prodromal females, and 13 prodromal males. We focused on high and moderate impact events because those are the most likely changes to result in functional and/or structural consequences for relevant protein-coding sequences. High impact editing events include those A-to-G or T-to-C nucleotide changes that result in large duplications or deletions, deleted or duplicated exons, frameshifts, gene deletions, etc. Moderate impact editing events include those that result in exon deletion or duplication, codon deletion or insertion, nonsynonymous coding, etc. [73]. Additionally, we compared the proportion of high and moderate impact editing events occurring in NCBI PD genes independently, between sample groups using Chi-square test.

We calculated editing index using BAM files from PPMI (hg19) remapped to hg37. Similar to Roth et al., 2019[83] approach, we defined editing index as the ratio of the number of A to G or T to C putative ADAR editing sites within sample data to the total number of A to G or T to C nucleotides mapped within samples. To create a standardized editing index per sample, we scaled the number of putative ADAR edits by 1,000,000 prior to the calculation of the indices. Variant calling was performed with GATK[73] using best practices described by Plonski et al. [69] through which only variants for sites with MAPQ ≥ 40 are called.

Differential expression was analyzed via DESEQ2[84] R package version 3.18 (R4.3.2)[85] with default parameters and results filtered for an adjusted p-value < 0.05 and log2 Fold Change less than −0.58 or greater than 0.58. The latter cut-off allowed us to consider genes that exhibited at least 50% change in their expression in either direction. Gene lists with greater than 2 members identified as being differentially expressed were analyzed via Reactome[79], or in the case of single genes, analyzed for function and potential connections to disease status. Gene lists were compared to the NCBI PD gene list[75] to identify differentially expressed genes that may be more likely to contribute to PD physiology. When Reactome[79] was used for pathway analysis for gene lists other than the NCBI PD gene list, filters of FDR <0.05 and at least 10% of entities in pathways were used. ADAR isoform expression was also analyzed with DESEQ2 R package version 2.18 (R4.3.2)[84] with read counts of less than 10 disregarded.

Chi-square test was used to compare the numbers of editing events across various categories, such as editing events with high and moderate impact, protein coding, and nonsense-mediated decay (NMD), using GraphPad Prism (version 10.1.2)[86] when categorical data was being analyzed. For ADAR expression, non-parametric Kruskal-Wallis and Mann-Whitney tests were used to compare PD patients, healthy patients, and prodromal patients, and pairwise comparisons of PD subsets, respectively. For comparing the total number of putative ADAR editing events per sample, as a proxy of ADAR editing levels, non-parametric Mann-Whitney test was used. For comparisons of ADAR isoform, unpaired t-test was used.

## 3 Results

### 3.1 Assessment of ADAR editing

To clarify, this assessment of ADAR editing in PD is not meant to infer that putative edits are “causative” to PD, but rather that they may “contribute” to the pathophysiology of PD. When total number of putative ADAR edits are considered, the highest average number of edits per sample was observed in prodromal females (120,281 ± 6,219 edits per sample), followed by PD males (119,971.2 ± 2,851 edits per sample), prodromal males (119,255.4 ± 5,028 edits per sample), healthy control females (118,694.7 ± 6,407 edits per sample), PD females (117,858.3 edits ± 4,444 per sample, and healthy control males (110,390.8 ± 4,345 edits per sample). When males and females are combined, prodromal samples have the highest number of putative ADAR edits with 119,601.9 ± 3,912 edits per sample followed by the PD group with 119,203.9 ± 2,423 edits per sample, and then healthy controls with 113,115.5 ± 3,602 edits per sample.

When range (difference between the highest number of editing events observed in a sample group and the lowest number of editing events observed in the group) is considered for major comparison groups, the largest range is observed in PD (range=163,348 events) followed by prodromal samples (range=160,243 editing events), and healthy controls (range= 116,762 editing events). When subgroups varying by sex are considered, the greatest range of ADAR editing events is observed in PD females (range = 163,348 editing events), prodromal males (range= 160,243 editing events), PD males (range = 139,562 editing events), prodromal females (range = 129,648 editing events), healthy control males (range = 112,347 editing events), and healthy control females (range = 100,143 editing events). There were no significant differences observed between sample groups when the number of putative ADAR edits are considered (Mann-Whitney, p> 0.05) (Supplementary Table 1).

**Table 1:** Mean number of ADAR editing events was calculated per sample for each group. ADAR editing events are those where A/G or U(T)/C edits are observed in the absence of a dbSNP[72] annotation. The highest mean number of ADAR editing events was observed in prodromal females (n=120,281 ± 6,219), followed by PD males (n=119,971 ± 2,851), prodromal samples (n=119,602 ± 3,912), prodromal males (n=119,255 ± 5,028), PD samples (n=119,204 ± 2,423), healthy control females (n=118,695 ± 6407), PD females (n=117,858 ± 4,444), healthy controls (n=113,116 ± 3,602), and healthy control males (n=110,391 ± 4,345). When editing index is calculated, which shows the proportion of A and T nucleotides in the raw RNA-seq data that represent putative ADAR edits, the highest mean editing index is observed in healthy control females (editing index =71.18 ± 2.803) followed by PD females (editing index = 69.56 ± 2.473), prodromal females (editing index = 68.33 ± 3.995), PD males (editing index = 66.56 ± 1.841), prodromal males (editing index = 64.50 ± 3.020) and healthy control males (editing index = 61.90 ± 3.088) however when the sexes are combined, PD samples had the highest editing index (67.65 ± 1.477) followed by prodromal samples (editing index = 65.79 ± 2.406) and healthy controls (editing index = 64.94 ± 2.322).

| Group | Mean number of ADAR editing events (SEM) per sample for each group | Mean Editing Index (SEM) per sample for each group |
| --- | --- | --- |
| Healthy Control Males (n=43) | 110,391 ± 4,345 | 61.90 ± 3.088 |
| Healthy Control Females (n=21) | 118,695 ± 6,407 | 71.18 ± 2.803 |
| Healthy Controls (n=64) | 113,116 ± 3,602 | 64.94 ± 2.322 |
| PD Males (n=114) | 119,971 ± 2,851 | 66.56 ± 1.841 |
| PD Females (n=65) | 117,858 ± 4,444 | 69.56 ± 2.473 |
| PD (n=179) | 119,204 ± 2,423 | 67.65 ± 1.477 |
| Prodromal Males (n=49) | 119,255 ± 5,028 | 64.50 ± 3.020 |
| Prodromal Females (n=25) | 120,281 ± 6,219 | 68.33 ± 3.995 |
| Prodromal (n=74) | 119,602 ± 3,912 | 65.79 ± 2.406 |

When editing index is calculated, which reflects the proportion of edited sites among the total set of A and T nucleotides that is present in a particular sample, the highest mean editing index is observed in healthy control females (editing index = 71.18 ± 2.803) followed by PD females (editing index = 69.56 ± 2.473), prodromal females (editing index = 68.33 ± 3.995), PD males (editing index = 66.56 ± 1.841), prodromal males (editing index = 64.50 ± 3.020) and healthy control males (editing index = 61.90 ± 3.088) however when the sexes are combined, PD samples had the highest editing index (67.65 ± 1.477) followed by prodromal samples (editing index = 65.79 ± 2.406) and healthy controls (editing index = 64.94 ± 2.322). Range of editing index (difference between the highest editing index observed in a sample group and the lowest editing index observed in the group) shows a slightly different trend with the highest range observed in PD females (range = 88.54), healthy control males (range = 80.76), PD males (range = 76.92), prodromal males (range = 75.06), prodromal females (range = 65.18), and healthy control females (range = 53.69) when the sexes are separated, but when combined the largest range of editing index is observed in healthy controls (range = 90.81) followed by PD samples (range = 88.54), and prodromal samples with the lowest range of editing indices (range = 75.06). Results are shown in Table 1 and Figure 1.

### 3.2 The NCBI PD gene list is over-enriched with Immune and Cell Signaling pathways

A list of 801 unique genes with suspected associations to PD were downloaded from NCBI Gene[75] (downloaded November 4, 2023) and converted to Ensembl gene and transcript IDs, resulting in 901 gene and 7311 transcript IDs utilizing Ensembl Biomart[76]. Reactome[79] pathway analysis was performed utilizing two variations of stringency: one filtering for FDR < 0.05 and another filtering for p value < 0.05 with at least 50% of entities in the pathway as designated by Reactome[79]. With the most stringent protocols (FDR < 0.05), NCBI PD genes were overrepresented in Reactome pathways Interleukin-4 and Interleukin-13 signaling (27%), Neutrophil degranulation (10.7%), Interleukin-10 signaling (31.4%), Post-translational protein phosphorylation (18.4%) and Platelet degranulation (17.0%). With the less stringent protocol of p value < 0.05 and at least 50% of entities within the pathway represented as designated by Reactome, Deletions in the AXIN1 gene destabilize the destruction complex, TLR3 deficiency, Defective OGG1 (DNA glycosylase enzyme) localization, Loss of function of TP53 in Cancer, Loss of function of TP53 in cancer due to loss of tetramerization ability, and Defective OGG1 substrate binding were represented by 100% of the entities in the pathway. Other groups overrepresented are Dopamine receptors (83%), Defective Base Excision Repair Associated with OGG1 (67 %), Transfer of LPS from LBP carrier to CD14 (67%), SMAC (DIABLO)-mediated dissociation of IAP caspase complexes (57%), NTRK2 activates RAC1 (57%), NFE2L2 regulating inflammation associated genes (57%), and MECP2 regulates transcription of neuronal ligands (54%), NTF3 activates NTRK2 (TRKB) signaling (50%), and BDNF activates NTRK2 (TRKB) signaling (50%). For clarity, this gene list will be referred to as NCBI PD gene list throughout this analysis. Results of Reactome[79] analysis of the NCBI PD gene list are found in Figures 2 and 3.

### 3.3 Variations in high and moderate impact ADAR edits in comparison groups

SnpEff[74] annotations of putative ADAR editing events were filtered for genes in which high or moderate impact editing events occurred within genes from the NCBI PD Gene[75] List and genome-wide. Entries with high or moderate impact editing events by coordinate that were found in less than 25% of samples were removed to identify the following numbers of genes with high or moderate impact edits: 205 in healthy control males, 213 in healthy control females, 240 in healthy controls, 217 in PD males, 217 in PD females, 228 in PD samples, 210 in prodromal males, 218 in prodromal females, and 243 genes with high or moderate impact edits in all prodromal samples. Genes with high or moderate impact protein coding edits were observed in sample groups as follows: 151 genes in healthy control males, 157 in healthy control females, 175 in healthy controls, 158 in PD males, 160 in PD females, 167 in PD, 157 genes in prodromal males, 154 in prodromal females, and 176 genes in the prodromal group. High or moderate impact edits in PD genes were observed in healthy control males (n=6 different genes), healthy control females (n=8 different genes), healthy controls (n=8 different genes), PD males (n=8 different genes), PD females (n=9 different genes), PD samples (n=9 different genes), prodromal males (n=7 different genes), prodromal females (n=8 different genes), and prodromal samples (n=8 different genes). In PD genes, high or moderate protein coding edits were observed in 6 genes in healthy control males, 8 genes in healthy control females, 8 genes in Healthy Controls, 8 genes in PD males, 8 genes in PD females, 8 genes in PD, 7 genes in prodromal males, 7 genes in prodromal females, and 7 genes in prodromal samples.

When genes in which protein coding high or moderate edits were observed, gene lists differed slightly per group when all genes are considered however when only PD genes are analyzed, some comparisons such as when healthy controls are compared to PD, PD females are compared to PD males, and prodromal females are compared to prodromal males, show no difference in the gene lists in which high or moderate impact protein coding edits are found. Gene lists can be found in Supplementary File 2.

Putative ADAR edits occurring in at least 25% of samples in a group were analyzed via SnpEff[74] and filtered for edits occurring in at least 25% of samples. The number of high and moderate impact edits and associated biotypes genome-wide and in PD genes were compared between healthy controls and PD, healthy controls and prodromal, PD and prodromal, healthy control males and pd males, healthy control males and prodromal males, PD males and prodromal males, healthy control females and PD females, healthy control females and prodromal females, PD females and prodromal females, healthy control females vs healthy control males, PD females and PD males, and prodromal females and prodromal males. Comparison groups can be found in Supplementary Table 2.

**Table 2:** High or Moderate Impact NMD Edits in PD genes. Significance is reached when comparing NMD edits in PD genes between Healthy Controls to PD samples (Chi-square, p=0.0106, increased proportionately in PD), Healthy Controls to prodromal samples (Chi-square, p= 0.0063, Increased proportionately In prodromal), healthy control females and PD females (Chi-square, p=0.0106, Increased proportionately In PD females), healthy control females and prodromal females (Chi-square, p=0.0049, Increased proportionately In prodromal females), PD females and PD males (Chi-square, p=<0.0001, increased proportionately in PD females), and prodromal females and prodromal males (Chi-square, p=<0.0001, increased proportionately in prodromal females).

|  | Healthy Controls | PD | Prodromal | Healthy Control Males | Healthy Control Females | PD Males | PD Females | Prodromal Males | Prodromal Females |
| --- | --- | --- | --- | --- | --- | --- | --- | --- | --- |
| Healthy Controls (n=64) | ----- | $p=0.0106$<br>^PD | ----- | ----- | ----- | ----- | ----- | ----- | ----- |
| PD (n=179) | ----- | ----- | $p=.7417$ | ----- | ----- | ----- | ----- | ----- | ----- |
| Prodromal (n=74) | $p=0.0063$<br>^Prodromal | ----- | ----- | ----- | ----- | ----- | ----- | ----- | ----- |
| Healthy Control Males (n=43) | ----- | ----- | ----- | ----- | ----- | N/A | ----- | N/A | ----- |
| Healthy Control Females (n=21) | ----- | ----- | ----- | N/A | ----- | ----- | ----- | ----- | ----- |
| PD Males (n=114) | ----- | ----- | ----- | ----- | ----- | ----- | $p<0.0001$<br>^PD Females | ----- | ----- |
| PD Females (n=65) | ----- | ----- | ----- | ----- | $p=0.0106$<br>^PD Females | ----- | ----- | ----- | ----- |
| Prodromal Males (n=49) | ----- | ----- | ----- | ----- | ----- | N/A | ----- | ----- | $p<0.0001$<br>^Prodromal Females |
| Prodromal Females (n=25) | ----- | ----- | ----- | ----- | $p=0.0049$<br>^Prodromal Females | ----- | $p=.6359$ | ----- | ----- |

Difference in protein coding and nonsense-mediated decay biotypes with high or moderate impact were analyzed as these biotypes were consistently represented in sample groups. Significant differences were observed when comparing the number of high and moderate impact edits genome-wide between healthy control males and prodromal males (Chi-square, p = 0.0493, increased proportionately in healthy control males), healthy control males and healthy control females (Chi-square, p = 0.0018, increased proportionately in healthy control males), and neared significance when comparing healthy control males to PD males (Chi-square, p = 0.075, increased proportionately in healthy control males). When comparing the number of high or moderate impact protein coding edits genome-wide, significance was achieved when comparing healthy control females and healthy control males (Chi-square, p=0.0184, increased proportionately in healthy control males). When high or moderate impact edits genome-wide result in nonsense-mediated decay, significance is reached when comparing PD samples to prodromal samples (Chi-square, p=0.0050, increased proportionately in PD), and PD males and prodromal males (Chi-square, p=0.0083, increased proportionately in PD).

When only PD genes are considered, no significant difference is reached (Chi-square test) between comparison groups when all high or moderate impact edits are considered or when protein coding high or moderate impact edits are analyzed; however, significance is achieved when considering the number of high and moderate impact edits in PD genes resulting in NMD. Significance is reached when comparing NMD edits in PD genes between healthy controls to PD samples (Chi-square, p=0.0106, increased proportionately in PD), healthy controls to prodromal samples (Chi-square, p= 0.0063, increased proportionately in prodromal), healthy control females and PD females (Chi-square, p=0.0106, increased proportionately in PD females), healthy control females and prodromal females (Chi-square, p=0.0049, increased proportionately in prodromal females), PD females and PD males (Chi-square, p=<0.0001, increased proportionately in PD females), and prodromal females and prodromal males (Chi-square, p=<0.0001, increased proportionately in prodromal females). The number of different genes (genome-wide) and PD genes observed with putative ADAR edits resulting in high or moderate impact, high or moderate impact protein coding or NMD edits across sample groups can be found in Supplementary Table 3. Statistical results can be found in Table 2 and Supplementary Tables 4-8. Comparisons of editing impacts can be found in Figures 4 and 5 (a-e).

### 3.4 ADAR expression is highest in PD and prodromal groups

Differential gene expression analysis with DESeq2 did not identify ADAR genes as differentially expressed as per our filtering criteria (log_2_ Fold Change < −0.58 or > 0.58, padj=<0.05) when the entire transcriptomes of male and female healthy controls, PD, and prodromal patients were considered. However, because there is a nuanced non-linear relationship between the levels of ADAR editing and expression of individual ADAR genes[87], we wanted to explore whether any differences in normalized ADAR genes expression can be identified among subject groups. ADAR expression levels as transcripts per million (TPM) were compared between groups revealing marked differences between groups; however, no significance was achieved (Kruskal-Wallis test, p >0.05). The highest expression of ADAR (ADAR1), which includes the interferon-inducible isoform ADARp150[88], was seen in PD male samples with 49.88 mean TPM (±2.604) followed by PD females (mean TPM= 48.26 ±3.570) and prodromal females (mean TPM =47.84 ±5.067). The lowest expression of ADAR was demonstrated in prodromal males (mean TPM = 42.08 ±3.179) followed by healthy control females (mean TPM= 43.25 ±5.080) and healthy control males (mean TPM = 44.70 ±3.559).

When males and females are combined, the PD groups shows the highest expression of ADAR (mean TPM= 49.77±2.614) followed by healthy controls (mean TPM= 44.22±2.893) and prodromal samples (mean TPM= 44.02±2.712) (Fig 13). When range is considered for ADAR expression within a sample set, PD females show the highest range (min=6.905 TPM, max= 182.7 TPM, range = 175.8) followed by PD males (min=8.298, max = 147.6, range= 139.3), healthy control males (min=10.96 TPM, max = 122.7 TPM, range = 111.8), prodromal males (min=7.018 TPM, max = 108.7 TPM, range = 101.2), prodromal females (min=9.475 TPM, max = 106.3 TPM, range = 96.80), then healthy control females (min= 10.74 TPM, max=103.4 TPM, range = 92.63). When the sexes are combined, the PD group shows the highest range of ADAR expression (range = 175.8 TPM) followed by healthy controls (range =112 TPM), then prodromal samples (range = 101.2 TPM). Results are shown in Figure 6 (a,b).

The expression of ADARB1 (ADAR2) was similar between all groups ranging from a mean of 2.707±.2474 (prodromal males) to a mean of 3.291±.4080 (prodromal females) (Fig 12). When males and females are combined, ADAR2 expression is highest in PD (3.105 TPM±.1235) followed by healthy controls (2.969±.2077), and prodromal samples (2.904±.2149) (Fig 13). The range of ADAR2 expression was highest in PD males (min=.8416 TPM, max = 11.66, range = 10.82) followed by healthy control males (min=.8420 TPM, max=10.97 TPM, range = 10.12), prodromal females (min= .7783 TPM, max = 7.747 TPM, range = 6.969), prodromal males (min =.4978 TPM, max = 7.220 TPM, range = 6.4723), PD females (min= .5087 TPM, max = 6.910 TPM, range = 6.401, and healthy control females (min = 1.236 TPM, max = 6.059 TPM, range = 4.823). The PD group had the highest range of ADAR2 expression (range = 11.15) followed by the prodromal group (range = 7.249), and healthy control samples (range = 10.12) There was little expression of ADARB2 (ADAR3) in any group, as ADARB2 is generally limited to expression in the brain[89] however the highest expression was observed in PD (.8422±.09130), followed by prodromal samples (.8233±.1850) and healthy controls (.7171±.1677). Results are shown in Figure 7 (a,b).

The highest range of ADAR 3 expression was observed in prodromal males (range = 9.295 TPM), healthy control males (range = 8.631), PD males (range = 7.145 TPM), PD males (range = 7.145), PD females (range = 6.739), prodromal females (range = 2.497), and healthy control females (range = 1.57) with all groups having at least 1 sample with 0 expression. When the sexes are combined, prodromal samples show the highest range in expression of ADAR3 (range =9.295), followed by healthy controls (range = 8.631) and PD samples (range = 7.145). Results are shown in Figure 8 (a,b).

Differential expression of ADAR isoforms, p110 and p150, were also compared between major comparison groups showing no significant difference in expression (unpaired t-test, p>0.05). ADAR p150, induced by IFN, is primarily located in the cytoplasm while p110 is constitutively expressed and localizes in the nucleus[90]. When differential expression of ADAR p110 (ENST00000368471) was compared between healthy controls and PD, healthy controls and prodromal samples, and PD and prodromal samples, there was no significant difference in expression with p values of 0.07,0 .07, and 0.96 (unpaired t-test), respectively. Likewise, no significant differences in expression were observed for ADAR p150 (ENST00000368474) for comparisons between the major groups (healthy controls vs PD: p=0.48, healthy controls vs. prodromal: p=0.38, PD vs prodromal: p=0.8). Results are shown in Figures 9-11.

### 3.5 Differentially expressed genes between comparison groups

Five genes were differentially expressed when comparing PD and healthy controls. ENSG00000243695 (RP11-290L7.3), a TGF Beta-Inducible Nuclear Protein 1 Pseudogene (log_2_ Fold Change = - 1.31, padj=.03), ENSG00000265037 (MIR4707) (log_2_ Fold Change = −2.3, padj = 1.39 x 10_-8_) a microRNA involved in post-transcriptional gene regulation, ENSG00000108825 (PTGES3L-AARSD1) (log_2_ Fold Change = −2.4, padj = 1.04 x 10^−7^), a protein coding gene affiliated with Distal Spinal Muscular Atrophy[91], and ENSG00000200983 (SNORA1) (log_2_ Fold Change = −1.7, padj = 6.25 x 10^−5^), a small nucleolar RNA, were downregulated in controls. No Reactome pathways were identified as over-represented when analyzing these four genes. Only 1 gene from the NCBI PD gene list was differentially expressed in this analysis with ENSG00000258761 (RP11-24J19.1), an RNA gene affiliated with lncRNA with its transcript antisense to SLC03A1 which enables sodium-independent transmembrane transporter activity[92], was observed to be upregulated (log_2_ Fold Change = 1.1, padj = .04) in healthy controls.

One hundred and sixty-five genes were observed to be differentially expressed when analyzing expression in prodromal samples and healthy controls with 34 downregulated genes (includes 1 NCBI PD gene) and 131 upregulated genes (includes 1 NCBI PD gene) in healthy controls. Reactome[79] analysis of the 131 upregulated genes revealed Peptide chain elongation, Nonsense Mediated Decay (NMD) independent of the Exon Junction Complex, Eukaryotic Translation Elongation, Formation of a pool of free 40S subunits, Eukaryotic Translation Termination, Selenocysteine synthesis, Viral mRNA Translation, Response of ElF2AK4 (GCN2) to amino acid deficiency, SRP-dependent co-translational protein targeting to membrane, GTP hydrolysis and joining of the 60S ribosomal subunit, L13a-mediated translational silencing of Ceruloplasmin expression, Nonsense-mediated decay, NMD enhanced by the Exon Junction Complex (EJC), Eukaryotic Translation Initiation, Cap-dependent Translation Initiation, SARS-CoV-1 modulates host translation machinery, and Formation of the ternary complex (and subsequently 43S complex) as pathways over-represented by the gene list when filters of FDR <0.05 and containing at least 10% of entities in the pathway are utilized as described by Reactome[79]. One of the 131 genes also found on the NCBI PD gene list, RP11-529H20.5, is an alias of ATXN3, a deubiquitinating enzyme[93,94] with associations to spinocerebellar ataxia type 3 (Machado-Joseph Disease), an autosomal dominant neurologic disorder[95]. Reactome over-representation analysis of the 34 downregulated genes revealed 5 pathways including Microtubule-dependent trafficking of connexons from Golgi to the plasma membrane, Erythrocytes take up oxygen and release carbon dioxide, Transport of connexons to the plasma membrane, Regulation of cortical dendrite branching, and Ca^2+^ activated K^+^ channels (FDR=<0.05, ≥10% entities in pathway). One gene among the 34 genes with decreased expression in prodromal samples compared to healthy controls, HBB encoding hemoglobin subunit beta, which is related to sickle cell anemia and Beta-thalassemia, is also found on the NCBI PD gene list. Several studies have found HBB, among other blood-related genes, to be downregulated in PD[96–99]. Additionally, a study of differential gene expression in PPMI blood samples observed the downregulation of genes associated with erythrocyte development pathways[100].

Seventy-six genes are differentially expressed when assessing prodromal samples and PD samples with 53 upregulated genes (including 1 NCBI PD gene) and 23 downregulated genes (including 1 NCBI PD gene) in PD. Reactome[79] overrepresentation analysis of the 53 upregulated genes did not produce any pathways that met the filtering criteria (FDR <0.05 and 10% of entities in pathway) however, the list did include 1 gene also found on the NCBI PD gene list. RP11-529H20.5 (ENSG00000259634), alias ATXN3, an upregulated gene in the analysis of prodromal and control differential expression, (log_2_ Fold Change = .89, padj = .00067), was also upregulated when prodromal and PD samples are assessed (log_2_ Fold Change = .61, padj = .002). Reactome over-representation analysis of the 23 downregulated genes revealed 4 pathways that met the criteria FDR<0.05 and at least 10% entities in pathway: Microtubule-dependent trafficking of connexons from Golgi to the plasma membrane, Transport of connexons to the plasma membrane, Erythrocytes take up oxygen and release carbon dioxide, and Nectin/Neci trans heterodimerization. Once again, HBB gene was highlighted as a PD gene that was differentially expressed when analyzing prodromal samples relative to PD (log_2_ Fold Change = −1.97, padj = 4.66 x 10^−10^).

Six hundred and nine genes were observed to be differentially expressed between PD males and PD females with 55 genes upregulated (including 1 NCBI PD gene) and 554 genes downregulated (including 6 NCBI PD genes) in PD females. One Reactome over-representation pathway met the filtering criteria (FDR<0.05 and ≥10% entities in pathway) when analyzing the upregulated 55 genes: HDMs demethylate histones.. Of those 55 genes, Xist, is the lone differentially expressed gene from the NCBI PD gene list (log_2_ Fold Change = 8.90, padj = 1.97 x 10^−72^). Xist, coding for a lncRNA, is important for X inactivation in mammalian females with higher serum levels observed in both males and female PD patients and those levels associated with PD stage[101]. Xist has been suggested to play a role in the male bias in some disease processes, including neurodegenerative disorders like PD[102].

Reactome overrepresentations analysis of the 554 downregulated genes revealed no pathways that met the filtering criteria (FDR < 0.05, ≥ 10% entities in pathway). Six downregulated genes were included on the NCBI PD gene list. When the 6 PD genes are analyzed for pathway over-representation, Dopamine Receptors and Chylomicron clearance are highlighted with FDR <0.05 and ≥10% entities in pathway. As biological females typically do not have the SRY gene which is found on the Y chromosome, SRY gene is observed as downregulated (log_2_ Fold Change = −4.94, padj = 3.31 x 10^−25^) in females. And while upregulation of SRY in males may be expected, dysregulation of the SRY gene has been suggested to play a role in PD pathology with a recent study of rat brains and human neuroblastoma cell lines demonstrating persistent upregulation of SRY and that SRY inhibition may be a strategy for mitigating male PD progression[103].

NRXN2, encoding cell adhesion molecules and receptors in vertebrate nervous systems, is also downregulated (log_2_ Fold Change = -.93, padj = .008) in PD females. In a recent study in cell cultures, overexpression of NRXN2 led to elevated levels of proteins associated with AD, ALS, and PD[104]. MMP16, also a downregulated gene identified in this comparison (log_2_ Fold Change = −1.95, padj=.02), is a matrix metalloproteinase tasked with the breakdown of the extracellular matrix during embryonic development and tissue remodeling but has also been related to inflammation and metastasis[105,106]. Additionally, a variant in MMP16, rs60298754, was deemed significant in a GWAS study of variants related to PD[107]. GRIN2C, also downregulated (log_2_ Fold Change = -.78, padj = .005), encodes a subunit of N-methyl-D-aspartate (NMDA) receptor, a subtype of glutamate receptor in the CNS, and is related to memory, synaptic development and learning[91]. Pozdyshev et al.[108] (2022), in a recent PPMI brain sample study, identified differentially edited sites of which over half were ionotropic glutamate receptors. Other subunits of NMDA receptors such as GRIN1 and GRIN2D have been observed with elevated expression in PD patients compared to controls in a small PD differential expression study[109]. Zhang et al.[110] (2016) suggested that hyperactivity of NMDA receptors can lead to excitotoxicity and may play a role in neurodegenerative disorders like AD and PD. DRD4, is a subtype of the dopamine G-protein coupled receptor and is used as a target for medications used to treat schizophrenia and Parkinson’s Disease[91] and was observed as differentially expressed in this study (log_2_ Fold Change = -.97, padj = .002). Additionally, variants in DRD4 have been identified as a risk factor for PD diagnosis in a sample of patients from Eastern India[111] and further, DRD4 has been connected to the development of neuropsychiatric disorders such as attention deficit disorder and schizophrenia[112]. APOE, or Apolipoprotein E, functions to transport lipids in the plasma associating with chylomicrons to assist with their clearance from the serum[113] and is observed to be differentially expressed in this analysis (log_2_ Fold Change = −1.2, padj=.04). APOE ε4 allele is a risk factor for sporadic Alzheimer’s[114] with recent data suggesting that APOE ε4 plays a role in alpha-synuclein pathology[115,116] and current evidence inferring that the variant is associated with cognitive decline in PD[117–119].

Genes that were observed to be differentially edited in our previous work[82] were evaluated for differential expression although the prior analysis involved skeletal muscle samples rather than blood samples as in this study. ACTA1, coding for skeletal muscle protein actin, ARL6IP5, RPA1 family protein 3, and ATPA2, sarcoplasmic/endoplasmic reticulum calcium ATPase 2, did not meet the criteria used in this study for differential expression (log_2_ Fold Change = <-0.58 or >0.58 and padj<0.05).

## 4 Discussion

RNA editing is a phenomenon that adds dynamic regulation to gene expression and specifically, ADAR editing is a nuanced process that varies by individual, developmental stage, and possibly other unknown factors[87,120]. Because ADAR editing is non-linear, we cannot expect that patterns of ADAR editing will imitate patterns of ADAR expression however, we do observe that ADAR expression and the numbers of editing events putatively attributed to ADAR, follow similar patterns although without significance and that editing may be dysregulated in diseased samples.

When groups are separated by sex, the highest mean number of ADAR editing events per sample was observed in prodromal females, followed by PD males, prodromal males, healthy control females, PD females, and healthy control males. Within-group comparison by sex shows that there is a higher proportion of edits found in PD compared to healthy controls with no significance, but a slightly greater proportion of edits in prodromal compared to PD also with no significance (Mann Whitney, p>0,05). A higher proportion of edits are observed in PD males compared to PD females with no significance, however female samples contained more edits per sample when comparing healthy control and prodromal males and females (healthy control males vs healthy control females (Mann Whitney, p>0.05). When entire groups are considered, the highest mean number of ADAR editing events per sample was found in prodromal samples, followed by PD samples, and lastly, healthy controls.

While the findings that prodromal and PD samples contain the highest number of putative editing events is perhaps not surprising given the relationship between ADARp150 isoform, IFN induced expression, and the inflammatory aspect of PD physiology, there was no significant difference observed between comparison groups when ADAR expression in analyzed although ADAR1 expression was highest in PD samples when sexes are considered separately, and when combined (Kruskal-Wallis, p>0.05). It is interesting to note, that while ADAR1 expression is highest in PD males and females, the number of putative ADAR editing events is highest in prodromal samples when sexes are combined. ADAR2 and 3 expression was also highest in either prodromal or PD samples depending on whether sexes are combined. Other studies have observed decreased editing frequency [121] and significant differences in ADAR expression[122] in PPMI PD blood; however, these studies did not compare PD males to PD females or PD to prodromal in their analyses and included samples regardless of age; whereas, we chose samples from individuals 65 years or older.

Editing index was highest in PD samples and prodromal samples with the lowest mean editing index observed in healthy controls. When sexes are considered separately, the same trend is observed in males where PD males have the highest mean editing index followed by prodromal males and healthy control males; however, in females, a very different trend is observed. Mean editing index is highest in healthy control females followed by PD females and prodromal females. Editing index ranges are highest in healthy controls followed by PD samples and prodromal samples with males following the same pattern with healthy control males having the highest range followed by PD males and prodromal males; however, female editing range patterns were altered. Editing index range was highest in PD females, followed by prodromal females and healthy control females were the lowest. Our results suggest that the dysregulation of editing in the diseased state may be different between males and females.

SnpEff analysis of putative ADAR editing outcomes reveals significant differences when comparing the number of high and moderate impact NMD edits genome-wide and in PD genes when comparing PD to prodromal samples, healthy controls to PD and healthy controls to prodromal samples, with the higher proportion of NMD edits observed in PD when compared to either healthy controls or prodromal and in prodromal when compared to healthy controls.

When male and female healthy controls are compared within a group, significant differences in the number of high and moderate impact edits genome-wide and high and moderate impact protein coding edits are observed with a greater proportion of those edits occurring in healthy males in both cases. In PD, significance is observed when comparing the number of high and moderate impact NMD edits in PD genes although NMD edits are elevated in PD females and there are no NMD edits in PD genes in PD males. There is also a significantly higher proportion of high and moderate impact NMD edits in PD genes observed in prodromal females compared to prodromal males. Where significant differences were observed in within-group comparisons, male samples showed a greater proportion of the edits than females except in NMD edits in PD genes where PD females had more edits compared to PD males and in prodromal females when compared to prodromal males suggesting that the manner in which transcripts are edited may differ between the sexes.

Comparisons by sex across groups reveal significant differences in the number of high and moderate impact edits genome-wide when comparing healthy males to prodromal males which serves as the one time in this study when a healthy control group has a significantly higher proportion of edits than a diseased group (PD or prodromal) although significance is barely achieved. High and moderate impact edits genome-wide associated with NMD were significantly elevated in PD males when compared to prodromal males, in PD genes when comparing PD females to healthy control females, and in prodromal females when compared to healthy control females. These comparisons where significant differences were observed followed the same pattern as when the entire groups were compared in that a higher proportion of edits occurred in PD samples over prodromal or healthy controls.

In our previous analysis of ADAR editing in the skeletal muscle of PD patients, a significant difference in the number of high or moderate impact edits resulting in NMD was observed (Chi-square, p=0.0001) when comparing healthy age-matched males (n=9) to a small subset of PD male patients (n=4) who participated in exercise training although these results were obtained from a small sample[82]. In our current analysis, we also observe significant differences in NMD-associated edits between groups although exercise training was not a factor in this study. This editing pattern suggests differences in the outcomes of ADAR editing particularly in NMD when comparing healthy controls to diseased states (prodromal or PD) and between males and females. NMD is an important process in detecting deleterious nonsense sequences in mRNA transcripts and eliminating them from the cell with their regulation serving an important role in neurodevelopment, neural differentiation, and neural maturation[44]. RNA misprocessing, which includes dysregulation of NMD, plays an important role in neurodegenerative diseases such as ALS and fronto-temporal dementia[42,44,123,124]. The role of ADAR editing in NMD that may contribute to PD pathophysiology should be further analyzed with an in-depth look at the specific genes affected; however, that remains out of the scope of this study.

Certainly, the genes identified as differentially expressed between PD, healthy control, and prodromal samples and particularly those suggested to play a role in PD pathology warrant further analysis for editing patterns. The comparison of expression in PD genes between PD males and PD females, particularly concerning transcripts of DRD4, GRIN2C, MMP16, NRXN2, and APOE, should be further scrutinized for altered editing patterns. Additionally, RP11-529H20.5 [ATXN3] and HBB, which were observed as differentially expressed between prodromal samples and healthy controls and between prodromal and PD samples should be further analyzed for editing patterns that may be significant in PD pathology. Several of the Reactome[79] pathway analyses of differentially expressed genes highlighted overrepresented pathways that relate to NMD which parallels our findings of ADAR editing associated with NMD disproportionately between comparison groups altogether suggesting that the role of NMD and NMD-related pathways in PD pathology should be further investigated.

PD research utilizing PPMI data[125] has spawned numerous studies analyzing imaging[126–129], cerebrospinal fluid (CSF)[129–132], dopamine transporter single photon emission tomography (DAT SPECT)[133,134], motor symptoms[135–137], and cognitive impairment[138,139] data from study participants for possible associations with PD etiology; however few studies have included ADAR editing in their analyses of PD.

Li et al.[122], observed 17 A-t o-I editing sites in NCOR1, BST1, and KANSL1 genes with potential risk associations for PD in a study of 380 PD and 178 healthy controls samples from PPMI. Additionally, alterations in ADAR editing at 57 sites correlated with the progression of cognitive symptoms in PD patients[122]. In another study of PPMI whole blood transcriptomes, 72 ADAR editing events were observed that affected miRNA binding, 8 of which may alter the expression of many other genes[121]. Editing within seed regions of miRNA transcripts can potentially change editing targets and cause further dysregulation of gene expression[121]. ADAR editing has also been associated with AD and PD in a study of blood and brain samples from both AD and PPMI PD patients identifying 198 loci in protein coding regions, of which 35 are potentially related to neurodegeneration[140]. Our study of PPMI whole blood transcriptomes also notes differences in ADAR editing when comparing healthy controls to diseased samples when focusing on the downstream consequences of editing events and further distinguishes between editing patterns in prodromal vs PD samples and patterns in PD males vs PD females. Additionally, in our focus on editing impacts, we note that edits associated with NMD may be of particular interest in PD etiology especially concerning the sex differences associated with PD phenotypes[141–143].

The Imbalance of male and female samples in our comparison groups is a limitation of this study as the observed trends may be influenced more heavily by male samples when the sexes are combined. In addition, inferences of the downstream consequences of ADAR edits as analyzed by Snpeff may be underestimated. Lastly, we did not attempt to separate our PD samples by suspected “axis” or PD phenotype[53,54] which could potentially illuminate differences in ADAR editing between the groups.

## 5 Conclusions

In this analysis, as in our previous study[82], we observed higher numbers of putative ADAR edits in PD samples compared to healthy controls, in addition there were higher levels of expression of IFN-inducible, ADAR1, although without significance. Notably, the greatest variation within sample groups of ADAR1 expression was observed in PD females when sex was considered and in PD samples when the sexes are combined, supporting the hypothesis of dysregulation in the disease state and potentially according to sex. In addition, we observe significant differences in the number of high and moderate impact edits when the whole genome is considered and when focusing on genes with suspected associations to PD (NCBI PD gene list) and when biotype, specifically NMD, is considered with higher proportions of those edits occurring in PD samples when compared to healthy controls.

When prodromal samples are compared to healthy controls, we observe trends similar to PD vs healthy control comparisons suggesting that there may be degrees of dysregulation to ADAR editing as one progresses with PD. In our previous work analyzing a small cohort of PD patients, PD female samples included significantly higher proportions of high and moderate impact edits when compared to PD males[82]; however in the current analysis, we do not achieve significance or higher proportions of edits when comparing high and moderate impact edits in males and females regardless of biotype, but certainly, PD females show higher proportions of edits when high and moderate impact edits resulting in NMD are assessed.

To conclude, we observe differences in ADAR expression, number of putative ADAR edits, editing index, and the number of high and moderate impact edits between comparison groups and diseased samples (either PD or prodromal) which often show higher levels of ADAR expression and edits. PD males and females also differ in ADAR expression, number of edits, and the number of high and moderate impact edits with males exhibiting elevations compared to females in all three categories except in the circumstance of those edits associated with NMD. Likewise, differentially expressed genes between comparison groups were tied to NMD-related pathways. Together, these results suggest that the dysregulation of ADAR editing plays a role in PD physiology and progression, editing patterns differ between PD males and PD females, and ADAR editing associated with NMD and genes functioning in NMD-related pathways may be integral to PD pathophysiology.

## Figure legends

Figure 1: Editing index is the ratio of the number of A to G or T to C putative ADAR edits within sample data to the total number of A to G or T to C nucleotides within samples. When editing index is calculated, the highest mean editing index is observed in healthy control females (editing index =71.18 ± 2.803) followed by PD females (editing index = 69.56 ± 2.473), prodromal females (editing index = 68.33 ± 3.995), PD males (editing index = 66.56 ± 1.841), prodromal males (editing index = 64.50 ± 3.020) and healthy control males (editing index = 61.90 ± 3.088) however when the sexes are combined, PD samples had the highest editing index (67.65 ± 1.477) followed by prodromal samples (editing index = 65.79 ± 2.406) and healthy controls (editing index = 64.94 ± 2.322. Range of editing index shows a slightly different trend with the highest range observed in PD females (range = 88.54), healthy control males (range = 80.76), PD males (range = 76.92), prodromal males (range = 75.06), prodromal females (range = 65.18), and healthy control females (range = 53.69) when the sexes are separated, but when combined the largest range of editing index is observed in healthy controls (range = 90.81) followed by PD samples (range = 88.54), and prodromal samples with the lowest range of editing indices (range = 75.06).

Figure 2: The NCBI PD gene list (downloaded November 4, 2023) was overrepresented in Reactome pathways that varied with filter stringency. a) Interleukin-10 signaling pathway was enriched with 31.4% of Reactome (Fabregat, 2019) entities represented in the NCBI PD gene list, Interleukin-4 and 13 signaling (27% of entities represented), Post-translational protein phosphorylation (18.4% of entities represented), Platelet degranulation (17% of entities represented), and Neutrophil degranulation (10.7% of entities represented) when filtering for pathways with FDR < 0.05. b) NCBI PD genes were overrepresented in 16 Reactome (Fabregat, 2019) pathways (p < 0.05) with at least 50% of Reactome entities represented in the pathway including 6 pathways with 100% of entities in pathways found in the NCBI PD gene list.

Figure 3: There are significant differences observed when comparing the number of high or moderate impact edits between sample groups. a) There is a significant difference observed when comparing the number of high or moderate impact edits in PD genes resulting in NMD in healthy controls to PD samples (Chi-square, p=0.0106) with a higher proportion of edits occurring in PD samples and when comparing healthy controls to prodromal samples (Chi-square, p=0.0063) with a higher proportion in prodromal samples. Shown is the mean number of high or moderate impact edits resulting in NMD in PD genes (healthy controls=0, PD = .11, prodromal =.12 average edits per sample). b) When comparing the number of high or moderate impact edits genome-wide resulting in NMD between sample groups, there is a significant difference between PD and prodromal samples (Chi-square, p=0.0050). Shown is the mean number of high or moderate impact edits resulting in NMD in all major comparison groups (healthy controls = 25.9, PD = 27.84, prodromal = 25.88 average edits per sample. c) Significant differences are observed (Chi-square) when comparing the number of high or moderate impact edits in PD genes that results in NMD in healthy control females and PD females (Chi-square, p= 0.0106), healthy control females to prodromal females (Chi-square, p=0.0049), PD females to PD males (Chi-square, p=0.0001) and prodromal females to prodromal males (Chi-square, p=0.0001). Shown are the mean number of high or moderate impact NMD edits in PD genes per sample (healthy control females = 0, healthy control males = 0, PD females = .30, PD males = 0, prodromal females = .36, and prodromal males = 0). d) Significant difference observed (Chi-square) when comparing the number of high or moderate impact edits resulting in NMD genome-wide when comparing PD males to prodromal males (Chi-square, p=0.0083). Shown is the mean number of high or moderate impact NMD edits genome-wide by comparison group (healthy control females = 28.2, healthy control males =24.77, PD females = 28.57, PD males = 27.42, prodromal females = 27.88, and prodromal males = 24.86 edits per sample). e) Significant differences are observed (Chi-square) when comparing the number of high or moderate impact edits genome-wide in healthy control females to healthy control males (Chi-square, p=0.0018) and healthy control males to prodromal males (Chi-square, p= 0.0493). Shown is the mean number of high or moderate impact edits genome-wide by sample group (healthy control females = 336.43, healthy control males = 304.12, PD females = 326.78, PD males = 324.49, prodromal females = 335.40, prodromal males = 319.94 average edits per sample). f) Significant differences are observed (Chi-square) when comparing the number of high or moderate impact protein coding edits genome-wide in healthy control females and healthy control males (Chi-square, p=0.0184). Shown is the mean number of high or moderate impact protein coding edits per sample group (healthy control females = 292.2, healthy control males = 261.89, PD females = 281.5, PD males = 280.64, prodromal females = 288.84, prodromal males = 280.35 average edits per sample). *p = < 0.05, **p= <0.01, ***p=0<.001

Figure 4: ADAR isoform expression varied between comparison groups with no significance (Kruskal-Wallis, p=>0.05). a) The highest mean expression of ADAR was observed in PD males and the lowest expression was observed in prodromal males with no significance observed between groups (Kruskal-Wallis, p=0.6993). The highest range of ADAR expression between samples within a sample group was observed in PD females (range = 175.8 TPM) and the lowest range of ADAR expression was observed in healthy control females (range = 92.63 TPM). b) When male and female samples are combined, the highest mean expression of ADAR was observed in PD samples with the lowest expression observed in prodromal samples with no significance observed between groups (Kruskal-Wallis, p=0.2955). The highest range of ADAR expression was observed in PD (range = 175.8 TPM) followed by healthy controls (range = 112 TPM) and prodromal samples (range = 101.2 TPM). c) The highest mean expression of ADAR2 was observed in prodromal females, followed by healthy females, PD males, PD females, healthy males and the lowest expression was observed in prodromal males with no significance observed between groups (Kruskal-Wallis, p=0.3878). d) When male and female samples are combined, the highest mean expression of ADAR2 was observed in PD samples with the lowest expression observed in prodromal samples with no significance observed between groups (Kruskal-Wallis, p=0.2703). e) The highest mean expression of ADAR3 was observed in prodromal males, followed by healthy males, PD males, PD females, prodromal males and the lowest expression was observed in healthy males with no significance observed between groups (Kruskal-Wallis, p=0.3354). f) When male and female samples are combined, the highest mean expression of ADAR3 was observed in PD samples with the lowest expression observed in healthy control samples with no significance observed between groups (Kruskal-Wallis, p=0.3577). g) ADAR isoform expression is differentially expressed without significance (unpaired t-test, p=>0.05) when comparing PD samples to healthy controls. Interferon-inducible ADAR isoform p150 (ENST00000368474) is not expressed with significant difference between PD and healthy controls (unpaired t-test, p=0.48). h) ADARp110 (ENST00000368471) expression (right) is not significantly different between PD and healthy control samples (unpaired t-test, p=0.066). i) ADAR isoform expression is differentially expressed without significance (unpaired t-test, p=>0.05) when comparing prodromal samples to healthy controls. Interferon-inducible ADAR isoform p150 (ENST00000368474) is not expressed with significant difference between prodromal and healthy controls (unpaired t-test, p=0.38). j) ADARp110 (ENST00000368471) expression is not significantly different between the prodromal and control groups (unpaired t-test, p=0.067). k) ADAR isoform expression is differentially expressed without significance (unpaired t-test, p=>0.05) when comparing prodromal samples to PD samples. Interferon-inducible ADAR isoform p150 (ENST00000368474) is not expressed with significant difference between prodromal and PD samples (unpaired t-test, p=0.8). l) ADARp110 (ENST00000368471) expression is not significantly different between PD and prodromal groups (unpaired t-test, p=0.96).

## Supplementary Materials

**Supplementary Table 1: The total number of A/G and T/C edits with no dbSNP annotation (Sherry et al**., **2001) were compared between groups with no significance observed (Mann-Whitney, p = > 0.05)**.

**Supplementary Table 2: Comparisons made between sample groups**.

**Supplementary Table 3: The number of different genes (genome-wide) and PD genes in which A to G or T to C edits were found varied between comparison groups**.

**Supplementary Table 4: High or Moderate Impact Edits Genome-wide**.

**Supplementary Table 5: High or Moderate Impact Edits in PD genes**.

**Supplementary Table 6: High or Moderate Impact Protein Coding Edits Genome-wide**.

**Supplementary Table 7: High or Moderate Impact Edits associated with NMD Genome-wide**.

**Supplementary Table 8: High or Moderate Impact Protein Coding Edits in PD genes**.

**Supplementary File 1: NCBI PD Gene List**

**Supplementary File 2: Edited gene lists by comparison group**

## Author Contributions

Conceptualization, Heather Milliken Mercer and Helen Piontkivska

Methodology, Heather Milliken Mercer and Helen Piontkivska

Software, Heather Milliken Mercer and Helen Piontkivska

Validation, Helen Piontkivska, Ayesha Tariq, and Aiswarya Mukundan

Formal analysis, Heather Milliken Mercer and Helen Piontkivska

Investigation, Heather Milliken Mercer and Helen Piontkivska

Resources, Heather Milliken Mercer and Helen Piontkivska

Data curation, Heather Milliken Mercer and Helen Piontkivska

Writing—original draft preparation, Heather Milliken Mercer

Writing—review and editing, Helen Piontkivska, Ayesha Tariq, and Aiswarya Mukundan

Visualization, Heather Milliken Mercer

Supervision, Helen Piontkivska

Project administration, Heather Milliken Mercer and Helen Piontkivska

Funding acquisition, Helen Piontkivska

All authors have read and agreed to the published version of the manuscript.

## Institutional Review Board Statement

This study was reviewed by the Institutional Review Board of Kent State University and qualified under exemption 4.

## Informed Consent Statement

**Not applicable**

## Data Availability Statement

This analysis used transcriptome sequencing data, obtained from the Parkinson’s Progression Markers Initiative (PPMI) upon request, www.ppmi-info.org.

## Acknowledgments

We would like to thank the participants and the researchers involved in the Parkinson’s Progression Markers Initiative (PPMI) for sharing their data to enable this study. PPMI – a public-private partnership – is funded by the Michael J. Fox Foundation for Parkinson’s Research and funding partners, including 4D Pharma, Abbvie, AcureX, Allergan, Amathus Therapeutics, Aligning Science Across Parkinson’s, AskBio, Avid Radiopharmaceuticals, BIAL, BioArctic, Biogen, Biohaven, BioLegend, BlueRock Therapeutics, Bristol-Myers Squibb, Calico Labs, Capsida Biotherapeutics, Celgene, Cerevel Therapeutics, Coave Therapeutics, DaCapo Brainscience, Denali, Edmond J. Safra Foundation, Eli Lilly, Gain Therapeutics, GE HealthCare, Genentech, GSK, Golub Capital, Handl Therapeutics, Insitro, Jazz Pharmaceuticals, Johnson & Johnson Innovative Medicine, Lundbeck, Merck, Meso Scale Discovery, Mission Therapeutics, Neurocrine Biosciences, Neuron23, Neuropore, Pfizer, Piramal, Prevail Therapeutics, Roche, Sanofi, Servier, Sun Pharma Advanced Research Company, Takeda, Teva, UCB, Vanqua Bio, Verily, Voyager Therapeutics, the Weston Family Foundation and Yumanity Therapeutics.

