## Supplemental_Materials for "The role of ADAR editing and nonsense-mediated decay in Parkinson’s Disease": Supplementary Table 1.docx

**Supplementary Table 1: The total number of A to G and T to C edits with no dbSNP annotation (Sherry et al., 2001) were compared between groups with no significance observed (Mann-Whitney, p = > 0.05).**

|  | **Results of Comparison in number of A/G and T/C edits (Mann-Whitney)** |
| --- | --- |
| **Healthy Controls (n=64) vs PD (n=179)** | p=.2337 |
| **Healthy Controls (n=64) vs Prodromal (n=74)** | p=.2995 |
| **PD (n=179) vs Prodromal (n=74** | P=.6196 |
| **Healthy Control Males (n=43) vs PD Males (n=65)** | P=.0887 |
| **Healthy Control Males (n=43) vs Prodromal Males (n=49)** | P=.2449 |
| **PD Males (n=114) vs Prodromal Males (n=49)** | P=.9342 |
| **Healthy Control Females (n=21) vs PD Females (n=65)** | P=.8419 |
| **Healthy Control Females (n=21) vs Prodromal Females (n=25)** | P=.9652 |
| **PD Females (n=65) vs Prodromal Females (n=25)** | P=.6035 |
| **Healthy Control Females (n=21) vs Healthy Control Males (n=43)** | P=.2861 |
| **PD Females (n=65) vs PD Males (n=114)** | P=.5687 |
| **Prodromal Females (n=25) vs Prodromal Males (n=49)** | p=.9548 |
