## Supplemental_Materials for "The role of ADAR editing and nonsense-mediated decay in Parkinson’s Disease": Supplementary Table 2.docx

**Supplementary Table 2- Comparisons made between sample groups.** When editing patterns are compared between healthy controls and PD samples, the effects of PD are compared. Comparisons between PD and prodromal samples analyzes the effects of PD progression and when sex is indicated, the effects of either PD or the effects of PD progression are being analyzed between males and females.

Healthy	PD	Prodromal	Healthy	Healthy	PD	PD	Prodromal	Prodromal

Controls	Control	Control	Males	Females	Males	Females

Males	Females

Healthy Controls	---------	Effects of PD	--------	--------	--------	--------	--------	--------	--------

(n=64)

PD	--------	--------	Effects of PD	--------	--------	--------	--------	--------	--------

(n=179)	Progression

Prodromal	Effects of PD	--------	--------	--------	--------	--------	--------	--------	--------

(n=74)	Progression

Healthy Control Males	--------	--------	--------	--------	--------	Effects of PD	--------	Effects of PD	-------

(n=43)	in Males	Progression in

Males

Healthy Control Females	--------	--------	--------	Effects of	--------	--------	--------	--------	--------

(n=21)	Sex

PD Males	--------	--------	---------	--------	--------	--------	Effects of	--------	--------

(n=114)	Sex in PD

PD Females	--------	--------	--------	--------	Effects of PD	--------	--------	--------	--------

(n=65)

Prodromal Males	--------	--------	--------	--------	--------	Effects of PD	--------	--------	Effects of PD

(n=49)	Progression	Progression

and Sex

Prodromal Females	--------	--------	--------	--------	Effects of PD	--------	Effects of PD	--------	-------

(n=25)	Progression in Females	Progression in Females
