## Supplemental_Materials for "The role of ADAR editing and nonsense-mediated decay in Parkinson’s Disease": Supplementary Table 3.docx

**Supplementary Table 3: The number of different genes (genome-wide) and PD genes in which A to G or T to C edits were found varied between comparison groups.** In addition, the number of high and moderate impact edits genome-wide and in PD genes and those that occur in protein coding regions or result in nonsense-mediated decay also differed between groups. Chi-square results of comparisons can be found in Table 5 and Supplementary Tables 4-8.

|  | Genes with High or Moderate Impact Edits genome-wide | PD Genes with High or Moderate Impact Edits | High or Moderate Impact Edits genome-wide | High or Moderate Impact Protein Coding Edits genome-wide | High or Moderate Impact Edits in PD Genes | High or Moderate Impact Protein Coding Edits in PD Genes | High or Moderate Impact NMD Edits genome-wide | High or Moderate Impact NMD Edits in PD genes |
| --- | --- | --- | --- | --- | --- | --- | --- | --- |
| Healthy Control Males (n=43) | 205 | 6 | 13077 | 11259 | 213 | 213 | 1065 | 0 |
| Healthy Control Females (n=21) | 213 | 8 | 7065 | 6136 | 130 | 130 | 592 | 0 |
| Healthy Controls (n=64) | 240 | 8 | 20142 | 17395 | 343 | 343 | 1657 | 0 |
| PD Males (n=114) | 217 | 8 | 36992 | 31993 | 677 | 677 | 3126 | 0 |
| PD Females (n=65) | 217 | 9 | 21241 | 8300 | 419 | 400 | 1857 | 19 |
| PD (n=179) | 228 | 9 | 58233 | 40293 | 1096 | 1077 | 4983 | 19 |
| Prodromal Males (n=49) | 210 | 7 | 15677 | 13737 | 292 | 292 | 1218 | 0 |
| Prodromal Females (n=25) | 218 | 8 | 8385 | 7221 | 153 | 144 | 697 | 9 |
| Prodromal (n=74) | 243 | 8 | 24062 | 20958 | 445 | 436 | 1915 | 9 |
