## Supplemental_Materials for "The role of ADAR editing and nonsense-mediated decay in Parkinson’s Disease": Supplementary Table 4.docx

**Supplementary Table 4–** **High or Moderate Impact Edits Genome-wide. The proportion of high or moderate impact edits genome-wide varied between comparison groups (Chi-square, p<.05).**

Healthy	PD	Prodromal	Healthy	Healthy	PD	PD	Prodromal	Prodromal

Controls	Control	Control	Males	Females	Males	Females

Males	Females

Healthy Controls	---------	p=.6273	--------	--------	--------	--------	--------	--------	--------

(n=64)

PD	--------	--------	p=.6991	--------	--------	--------	--------	--------	--------

(n=179)

Prodromal	p=.4680	--------	--------	--------	--------	--------	--------	--------	--------

(n=74)

Healthy Control Males	--------	--------	--------	--------	--------	p=.0747	--------	p=.0493	--------

(n=43)	^Healthy Control Males

Healthy Control Females	--------	--------	--------	p=.0018	--------	--------	--------	--------	--------

(n=21)	^Healthy Control Males

PD Males	--------	--------	---------	--------	--------	--------	p=.4715	--------	--------

(n=114)

PD Females	--------	--------	--------	--------	p=.1122	--------	--------	--------	--------

(n=65)

Prodromal Males	--------	--------	--------	--------	--------	p=.5887	--------	--------	p=.9874

(n=49)

Prodromal Females	--------	--------	--------	--------	p=.1608	--------	p=.9486	--------	--------

(n=25)
