## Supplemental_Materials for "The role of ADAR editing and nonsense-mediated decay in Parkinson’s Disease": Supplementary Table 5.docx

**Supplementary Table 5 –** **High or Moderate Impact Edits in PD genes. The proportion of high or moderate impact edits in PD genes did not differ between comparison groups (Chi-square, p>0.05).**

Healthy	PD	Prodromal	Healthy	Healthy	PD	PD	Prodromal	Prodromal

Controls	Control	Control	Males	Females	Males	Females

Males	Females

Healthy Controls	---------	p=.1324	--------	--------	--------	--------	--------	--------	--------

(n=64)

PD	--------	--------	p=.7100	--------	--------	--------	--------	--------	--------

(n=179)

Prodromal	p=.3151	--------	--------	--------	--------	--------	--------	--------	--------

(n=74)

Healthy Control Males	--------	--------	--------	--------	--------	p=.2702	--------	p=.2957	--------

(n=43)

Healthy Control Females	--------	--------	--------	p=.6263	--------	--------	--------	--------	--------

(n=21)

PD Males	--------	--------	---------	--------	--------	--------	p=.2512	--------	--------

(n=114)

PD Females	--------	--------	--------	--------	p=.3013	--------	--------	--------	--------

(n=65)

Prodromal Males	--------	--------	--------	--------	--------	p=.9135	--------	--------	p=.9572

(n=49)

Prodromal Females	--------	--------	--------	--------	p=.7714	--------	p=.4646	--------	--------

(n=25)
