## Supplemental_Materials for "The role of ADAR editing and nonsense-mediated decay in Parkinson’s Disease": Supplementary Table 6.docx

**Supplementary Table 6 – High or Moderate Impact Protein Coding Edits Genome-Wide varied by proportion between comparison groups (Chi-square, p<0.05).**

Healthy	PD	Prodromal	Healthy	Healthy	PD	PD	Prodromal	Prodromal

Controls	Control	Control	Males	Females	Males	Females

Males	Females

Healthy Controls	---------	p=.6651	--------	--------	--------	--------	--------	--------	--------

(n=64)

PD	--------	--------	p=.5700	--------	--------	--------	--------	--------	--------

(n=179)

Prodromal	p=.8755	--------	--------	--------	--------	--------	--------	--------	--------

(n=74)

Healthy Control Males	--------	--------	--------	--------	--------	p=.2142	--------	p=.6573	--------

(n=43)

Healthy Control Females	--------	--------	--------	p=.0184	--------	--------	--------	--------	--------

(n=21)	^healthy control males

PD Males	--------	--------	---------	--------	--------	--------	p=.2780	--------	--------

(n=114)

PD Females	--------	--------	--------	--------	p=.3520	--------	--------	--------	--------

(n=65)

Prodromal Males	--------	--------	--------	--------	--------	p=.4347	--------	--------	p=.2256

(n=49)

Prodromal Females	--------	--------	--------	--------	p=.4154	--------	p=.9764	--------	--------

(n=25)
