## Supplemental_Materials for "The role of ADAR editing and nonsense-mediated decay in Parkinson’s Disease": Supplementary Table 7.docx

**Supplementary Table 7 –** **High or Moderate Impact Edits associated with NMD genome-wide varied by proportion between comparison groups (Chi-square, p<0.05).**

Healthy	PD	Prodromal	Healthy	Healthy	PD	PD	Prodromal	Prodromal

Controls	Control	Control	Males	Females	Males	Females

Males	Females

Healthy Controls	---------	p=.2115	--------	--------	--------	--------	--------	--------	--------

(n=64)

PD	--------	--------	p=.0500	--------	--------	--------	--------	--------	--------

(n=179)	^PD

Prodromal	p=.2327	--------	--------	--------	--------	--------	--------	--------	--------

(n=74)

Healthy Control Males	--------	--------	--------	--------	--------	p=.5950	--------	p=.0936	--------

(n=43)

Healthy Control Females	--------	--------	--------	p=.7325	--------	--------	--------	--------	--------

(n=21)

PD Males	--------	--------	---------	--------	--------	--------	p=.3426	--------	--------

(n=114)

PD Females	--------	--------	--------	--------	p=.1737	--------	--------	--------	--------

(n=65)

Prodromal Males	--------	--------	--------	--------	--------	p=.0083	--------	--------	p=.1560

(n=49)	^PD males

Prodromal Females	--------	--------	--------	--------	p=.7944	--------	p=.2639	--------	--------

(n=25)
