## Supplemental_Materials for "The role of ADAR editing and nonsense-mediated decay in Parkinson’s Disease": Supplementary Table 8.docx

**Supplementary Table 8 – High or Moderate Impact Protein Coding Edits in PD genes did not vary by proportion between comparison groups (Chi-square, p>0.05).**

Healthy	PD	Prodromal	Healthy	Healthy	PD	PD	Prodromal	Prodromal

Controls	Control	Control	Males	Females	Males	Females

Males	Females

Healthy Controls	---------	p=.2229	--------	--------	--------	--------	--------	--------	--------

(n=64)

PD	--------	--------	p=.6743	--------	--------	--------	--------	--------	--------

(n=179)

Prodromal	p=.4738	--------	--------	--------	--------	--------	--------	--------	--------

(n=74)

Healthy Control Males	--------	--------	--------	--------	--------	p=.2702	--------	p=.2957	--------

(n=43)

Healthy Control Females	--------	--------	--------	p=.6263	--------	--------	--------	--------	--------

(n=21)

PD Males	--------	--------	---------	--------	--------	--------	p=.6939	--------	--------

(n=114)

PD Females	--------	--------	--------	--------	p=.5708	--------	--------	--------	--------

(n=65)

Prodromal Males	--------	--------	--------	--------	--------	p=.9135	--------	--------	p=.5162

(n=49)

Prodromal Females	--------	--------	--------	--------	p=.8293	--------	p=.3912	--------	--------

(n=25)
